## Supplemental text and figures for "CASPULE: A computational tool to study sticker spacer polymer condensates"

**Effect of timestep and viscosity on equilibrium and kinetic properties of condensates**

We consider an energy scale (Es = 6kT, Ens = 0.3kT) where the system remains as a single large cluster (Fig. 4). We simulate the system for a given amount of elapsed or simulation time (50 * $10^6$), but with different timesteps such that $t_{sim} = N_{step}\ \mathrm{d}t$. For instance, a timestep $(\mathrm{d}t)$ with 1 and 5 femtosecond needs 50 and 10 million steps $(N_{step})$, respectively, to reach identical elapsed time.

In LAMMPS (see documentation), a particle under the Langevin dynamics experiences a summation of three force terms: $F = F_{conservative} + F_{viscous} + F_{random}$. $F_{conservative}$ is computed from the inter-particle potential. $F_{viscous}$ is the viscous damping term that is proportional to the mass and velocity of the particle, $F_{viscous} = -\frac{m}{damp}v$ where "damp" is a time parameter that inversely sets the viscosity of the medium. $F_{random} \propto \sqrt{\frac{k_B T m}{dt\ damp}}$ is the random noise term to mimic the solvent effect implicitly.

Figure S1 summarizes the energetics of the system at $t_{damp} = 200$. We simulate the system with 8 different timesteps. As a rule of thumb, a smaller timestep usually yields more accurate results for numerical integration. Fig. S1A shows the potential energy (PE) trend of the system. Being the smallest, timestep = 1fs may serve as the ground truth. We notice that the PE remains invariant up to timestep = 10fs. It shows slight upward trend with timestep = 20 and beyond. As a useful control, we also display the kinetic energy (Fig. S1B) of the system which should not depend on the timestep. The deviation in PE mainly comes from the reduced degree of sticker bonding as captured in Fig. S1C. Larger timesteps cause more displacements which are likely to break more bonds, on average. However, in numerical simulations, there is always a trade-off between accuracy and computational cost. Fig. S1D shows the trend of execution time (CPU time). While 1fs timestep provides accurate results, it almost takes ~10x more time to simulate identical

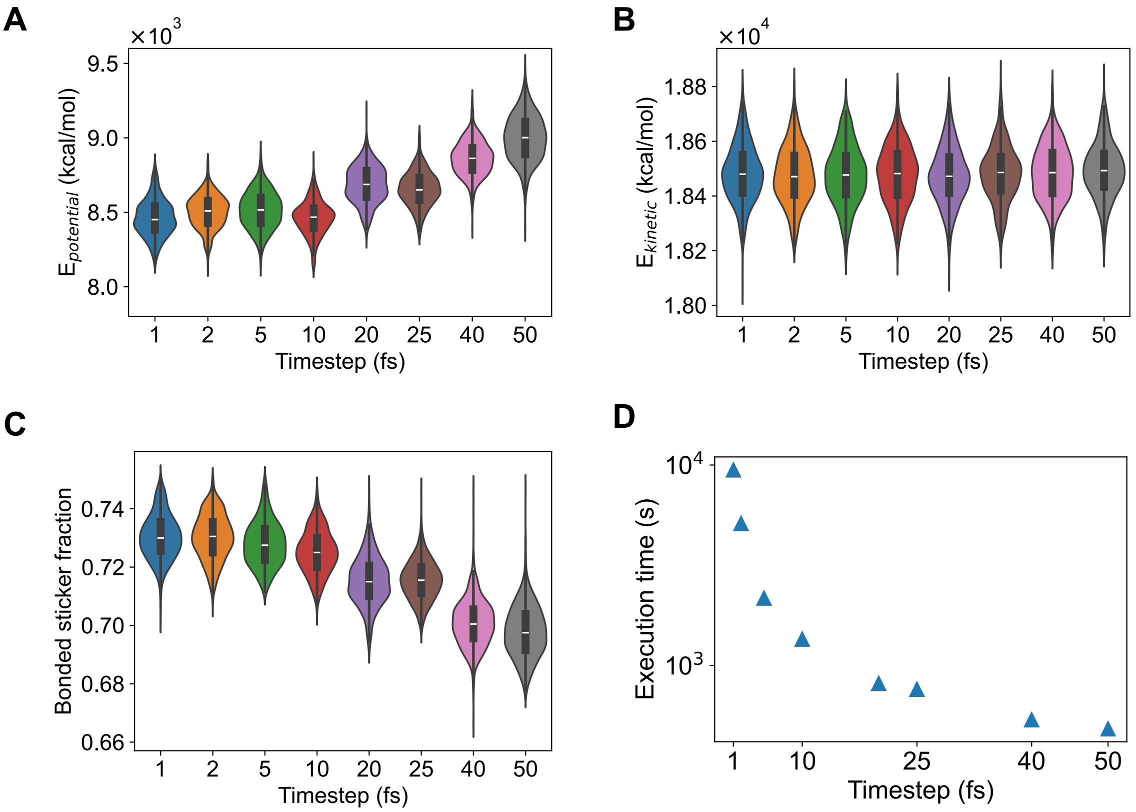


**Figure S1: Effect of timestep.** Trend of (A) potential energy (B) Kinetic energy (C) Sticker bonding statistics and (D) execution time. Energy parameter, Es = 6kT, Ens = 0.3kT. Damping parameter = 200 fs. Total elapsed or simulation time = 50 * $10^6$.

elapsed time compared to 10fs timestep. Fig. S2 extends the analysis for two another damping factor. For a smaller damp time ($t_{damp} = 200$), that signifies higher solvent viscosity, PE can tolerate larger timesteps and yield similar PE statistics. For a larger damping, the tolerance slightly goes down. In summary, for a medium with $t_{damp} = 500$, a timestep of 10 to 25 fs provides an optimal tradeoff in accuracy vs. cost space.

As mentioned earlier, the damping parameter can be thought to be inversely related to the medium viscosity. Since viscosity is an important kinetic parameter relevant in the study of kinetic arrest and condensate size distribution, we test the cluster motion with a series of damping parameters (Fig. S3). For a given damping time, we compute the mean squared displacement (MSD) of the cluster center and extract the diffusion coefficient (D) from the following relation: MSD = 6Dt (Fig. S3A). To our satisfaction, D scales linearly with $t_{damp}$ which implies $D \propto t_{damp} \Rightarrow D \propto 1/\eta$ where

$\eta$ is solvent viscosity. This scaling is consistent with Stokes-Einstein relation connecting the diffusivity of an object and solvent viscosity.

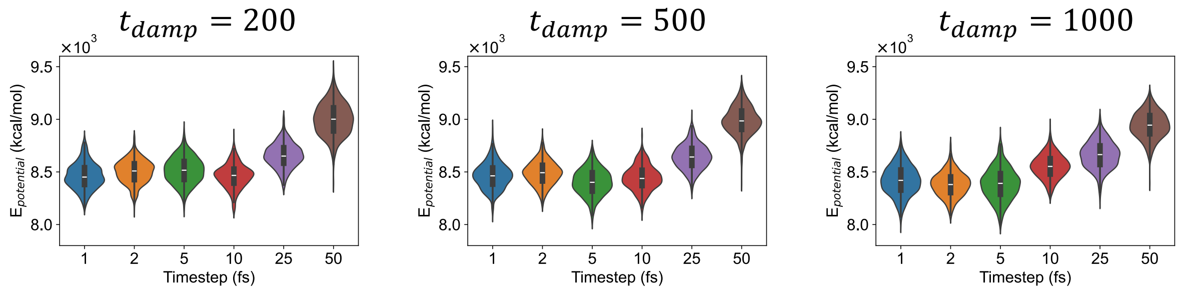


**Figure S2: Effect of damping time.** Trend of the potential energy at three different values of damping time. Energy parameter, Es = 6kT, Ens = 0.3kT. Damping parameter = 200 fs. Total elapsed or simulation time = 50 * $10^6$.

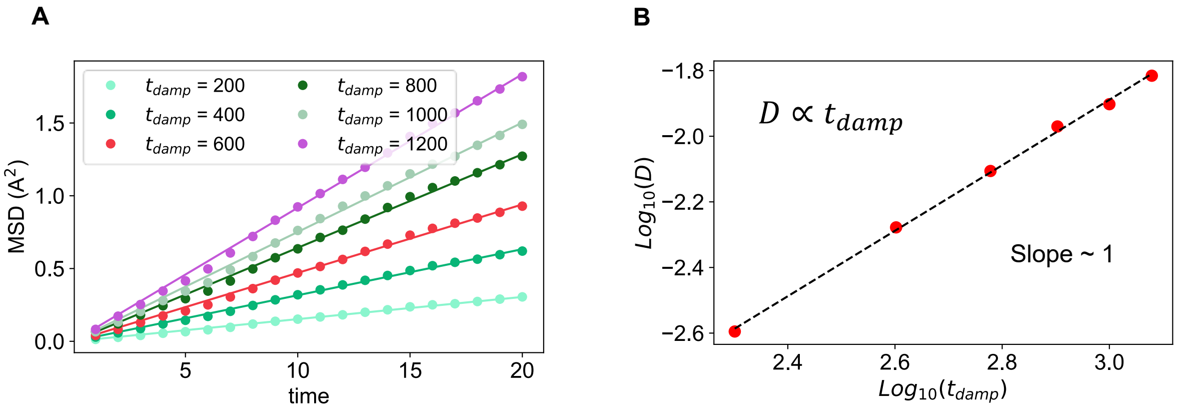


**Figure S3: Effect of damping time on cluster diffusion.** (A) Mean squared displacement vs. elapsed time plot for six different damping time (viscosity). The circles indicate actual displacements, and solid lines show the linear fit. Diffusion coefficient, D, is extracted from the linear fit: MSD = 6Dt (B) D vs. damping time in log-log scale. The red points are extracted Ds from MSD trajectories. The black dashed line is the linear fit which yields a slope ~ 1. In these simulations, timestep = 10 fs. Energy parameters, Es = 6kT, Ens = 0.3kT.
